## Supplements for "Activation-inducible CAR expression enables precise control over engineered CAR T cell function"

### Slide 1
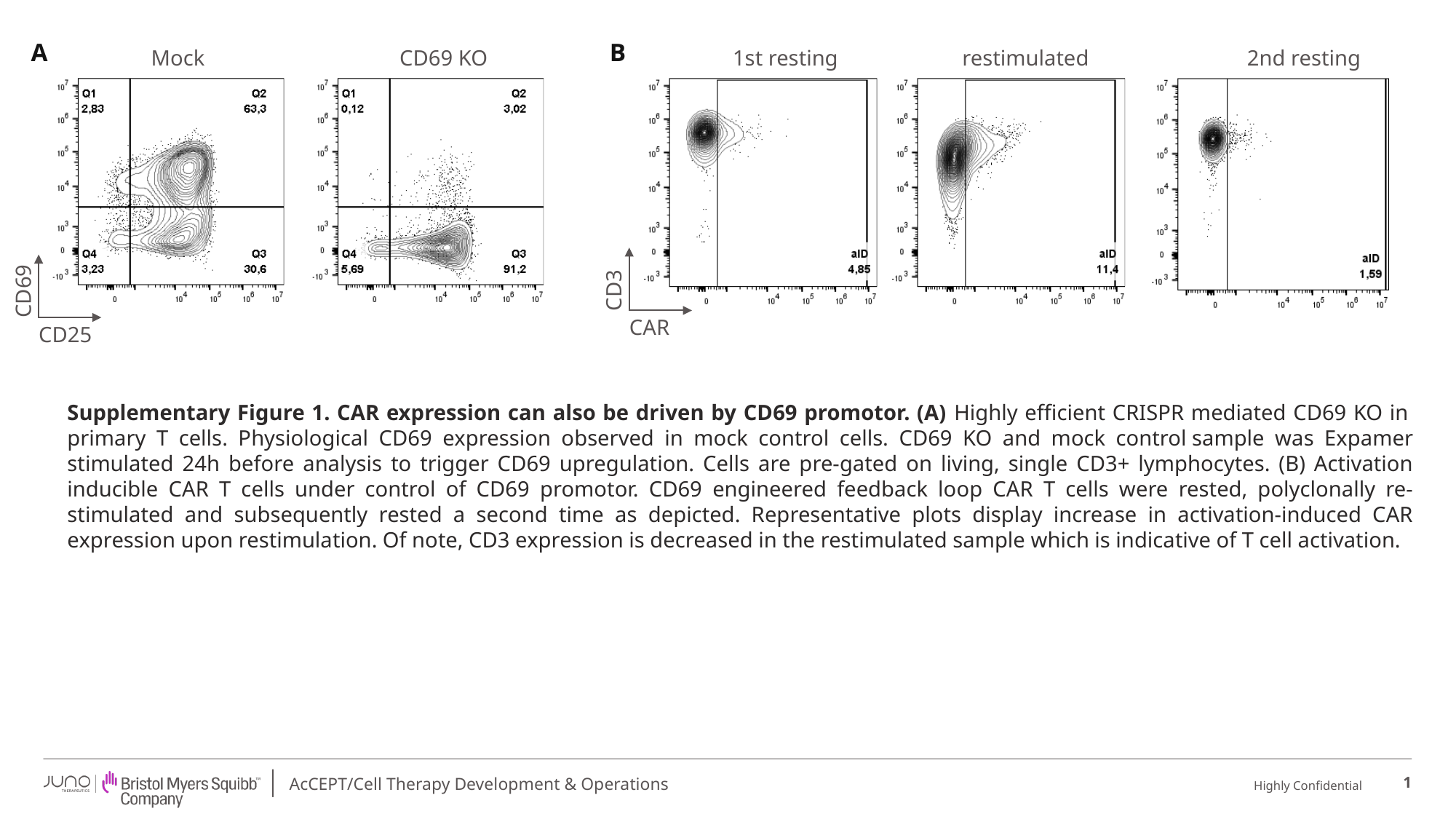

A
B
Mock
CD69 KO
1st resting
restimulated
2nd resting
CD3
CD69
CAR
CD25
Supplementary Figure 1. CAR expression can also be driven by CD69 promotor. (A) Highly efficient CRISPR mediated CD69 KO in  primary T cells. Physiological CD69 expression observed in mock control cells. CD69 KO and mock control sample was Expamer stimulated 24h before analysis to trigger CD69 upregulation. Cells are pre-gated on living, single CD3+ lymphocytes. (B) Activation inducible CAR T cells under control of CD69 promotor. CD69 engineered feedback loop CAR T cells were rested, polyclonally re-stimulated and subsequently rested a second time as depicted. Representative plots display increase in activation-induced CAR expression upon restimulation. Of note, CD3 expression is decreased in the restimulated sample which is indicative of T cell activation.
1

### Slide 2
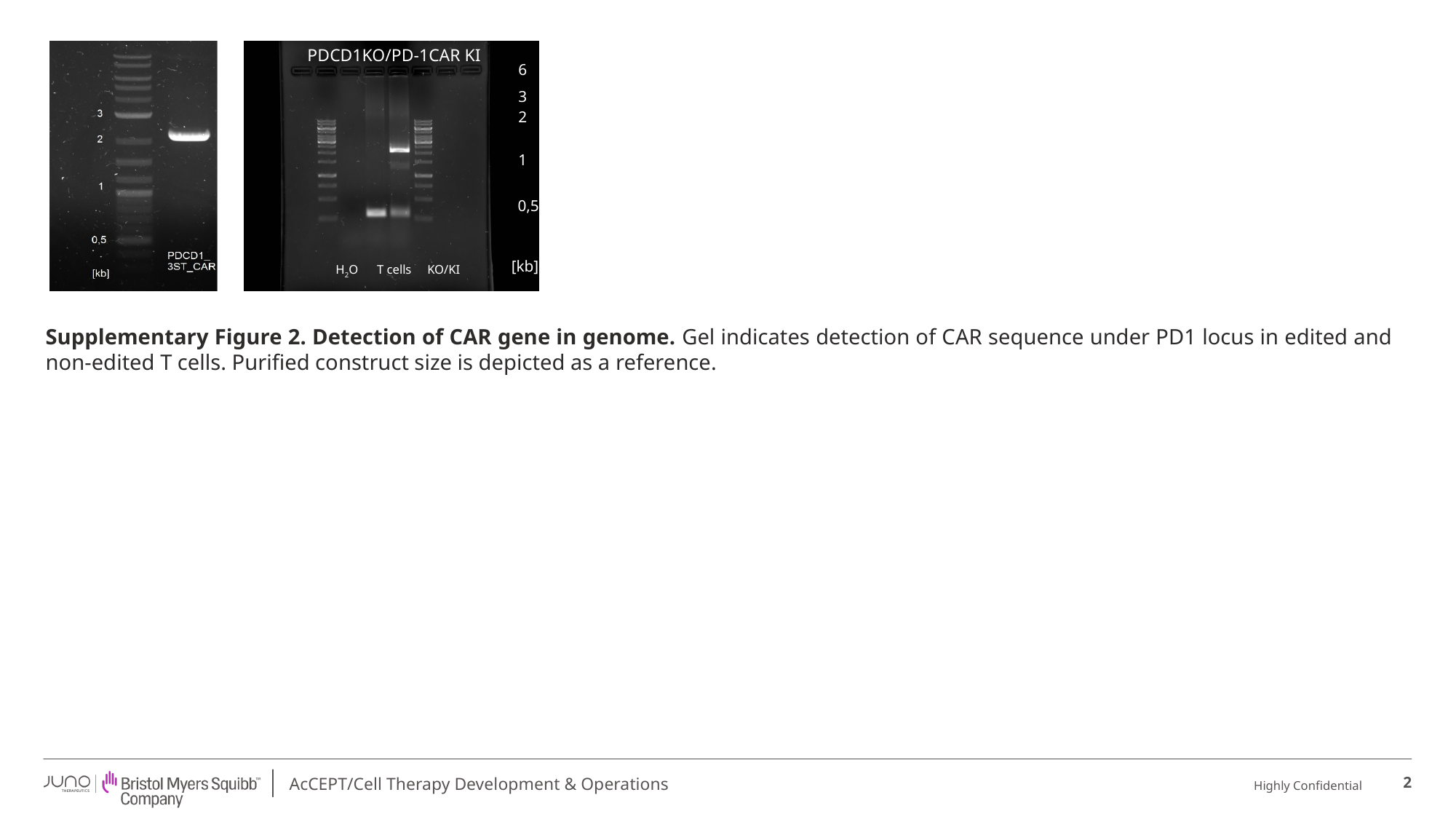

PDCD1KO/PD-1CAR KI
H_2_O
T cells
KO/KI
6
PDCD1KO/PD-1CAR KI
3
2
1
0,5
[kb]
H_2_O
T cells
KO/KI
Supplementary Figure 2. Detection of CAR gene in genome. Gel indicates detection of CAR sequence under PD1 locus in edited and non-edited T cells. Purified construct size is depicted as a reference.
2
